# An efficient gene targeting system using Δ*ku80* and functional analysis of Cyp51A in *Trichophyton rubrum*

**DOI:** 10.1101/2024.04.18.590175

**Authors:** Masaki Ishii, Tsuyoshi Yamada, Shinya Ohata

## Abstract

*Trichophyton rubrum* is one of the most frequently isolated fungi in patients with dermatophytosis. Despite its clinical significance, the molecular mechanisms of drug resistance and pathogenicity of *T. rubrum* remain to be elucidated because of the lack of genetic tools, such as efficient gene targeting systems. In this study, we generated a *T. rubrum* strain that lacks the nonhomologous end-joining-related gene *ku80* (Δ*ku80*) and then developed a highly efficient genetic recombination system with gene targeting efficiency that was 46 times higher than that using the wild-type strain. Cyp51A and Cyp51B are 14-α-lanosterol demethylase isozymes in *T. rubrum* that promote ergosterol biosynthesis and are the targets of azole antifungal drugs. The expression of *cyp51A* mRNA was induced by the addition of the azole antifungal drug efinaconazole, whereas no such induction was detected for *cyp51B*, suggesting that Cyp51A functions as an azole-responsive Cyp51 isozyme. To explore the contribution of Cyp51A to susceptibility to azole drugs, the neomycin phosphotransferase (*nptII*) gene cassette was inserted into the *cyp51A* 3′-UTR region of Δ*ku80* to destabilize the mRNA of *cyp51A*. In this mutant, although the expression level of *cyp51A* mRNA was comparable to that of the parent strain, the induction of *cyp51A* mRNA expression by efinaconazole was diminished. The minimum inhibitory concentration for several azole drugs of this strain was reduced, suggesting that dermatophyte Cyp51A contributes to the tolerance for azole drugs. These findings suggest that an efficient gene targeting system using Δ*ku80* in *T. rubrum* is applicable for analyzing genes encoding drug targets.

**Key Points:**

1. A novel gene targeting system using Δ*ku80* strain was established in *T. rubrum*
2. Cyp51A in *T. rubrum* responds to the azole antifungal drug efinaconazole
3. Cyp51A contributes to azole drug tolerance in *T. rubrum*

## INTRODUCTION

Dermatophytosis is a superficial fungal infection with symptoms such as itching, redness, and nail abnormalities. Tinea pedis (athlete’s foot), a type of dermatophytosis, affects approximately 10% of the world’s population (1). *Trichophyton rubrum*, the most common dermatophyte (2), is a clinically important organism that reduces the quality of life and has a unique life cycle as an anthropophilic dermatophyte that specifically inhabits human surface tissues. A limited class of antifungals, such as azole antifungals, are used in dermatophytosis treatment. Although drug resistance issues in *T. rubrum* have resulted in a need to elucidate the detailed molecular mechanisms of its drug resistance and identify and analyze drug targets (3, 4), these issues have not been completely clarified because of the underdevelopment of genetic methods in *T. rubrum*.

Homologous recombination (HR), a repair mechanism for DNA double-strand, is one of the most commonly used genetic engineering methods (5). This technique allows the precise insertion of any DNA fragment into the desired genomic region based on sequence homology. Nevertheless, eukaryotes also possess a nonhomologous end-joining (NHEJ) repair mechanism for double-strand breaks, which competes with HR-mediated insertion of DNA into target regions (6). To efficiently promote targeted integration via HR, several fungal species have been engineered by disrupting either of the Ku70/Ku80 complexes involved in NHEJ (7, 8). These strains have demonstrated the effectiveness of improving HR efficiency in various fungi (7, 8).

In this study, we established a highly efficient HR system using a *ku80*-deficient strain of *T. rubrum*. Using this established system, we developed a mutant in which the neomycin phosphotransferase (*nptII*) gene was inserted into the 3L-UTR region of *cyp51A*, which encodes a target for azole antifungals. When the azole antifungal drug efinaconazole was added, the magnitude of increase in *cyp51A* expression decreased in this mutant, which also exhibited sensitivity to ravuconazole and efinaconazole. This study would accelerate the production of genetically engineered strains to investigate the pathogenicity and drug resistance of *T. rubrum* and provide novel insights into antifungal targets.

## MATERIALS AND METHODS

### Fungal and bacterial strains and culture conditions

*T. rubrum* CBS118892 (9), a clinically isolated strain from a patient’s nail, was cultured on Sabouraud dextrose agar (SDA; 1% Bacto peptone, 4% glucose, 1.5% agar, pH unadjusted) at 28°C. The conidia of *T. rubrum* were prepared as described previously (10). We confirmed the sequence of *cyp51A* and *cyp51B* as well as their promoters and terminators.

### Plasmid construction

To construct a *ku80*-targeting vector, pAg1-Δ*ku80-flp*, approximately 2.1 and 1.5Lkb of the 5L- and 3L-UTR fragments, respectively, of the *ku80* open reading frame (ORF) were amplified from *T. rubrum* genomic DNA by polymerase chain reaction (PCR). The PCR products of the 5L- and 3L-UTR fragments were cleaved by *Spe*I/*Apa*I and *Bgl*II/*Kpn*I, respectively. The plasmid backbone of pAg1 and the FLP/FRT module of pMRV-TmKu80/T2 were cleaved by *Spe*I/*Kpn*I and *Apa*I/*Bam*HI, respectively (11). These fragments were joined using Ligation high version 2 (TOYOBO, Japan). To construct a *cyp51A* 3L-UTR-targeting vector, pAg1-*cyp51A*-3L-UTR, the *cyp51A* ORF and 1.5 kb of the 3L-UTR fragment of *cyp51* ORF were amplified from *T. rubrum* genomic DNA by PCR. The *neomycin phosphotransferase* gene cassette, which consists of *E. coli* neomycin phosphotransferase (*nptII*), *Aspergillus nidulans trpC* promoter (*PtrpC*), and *Aspergillus fumigatus cgrA* terminator (*TcgrA*), was cleaved from pMRV-TmKu80/T2 using *Apa*I and *Cla*I. These fragments were joined using an In-Fusion HD Cloning Kit (TaKaRa Bio, Japan). The primers used in this study are shown in Table 1.

**Table 1.**
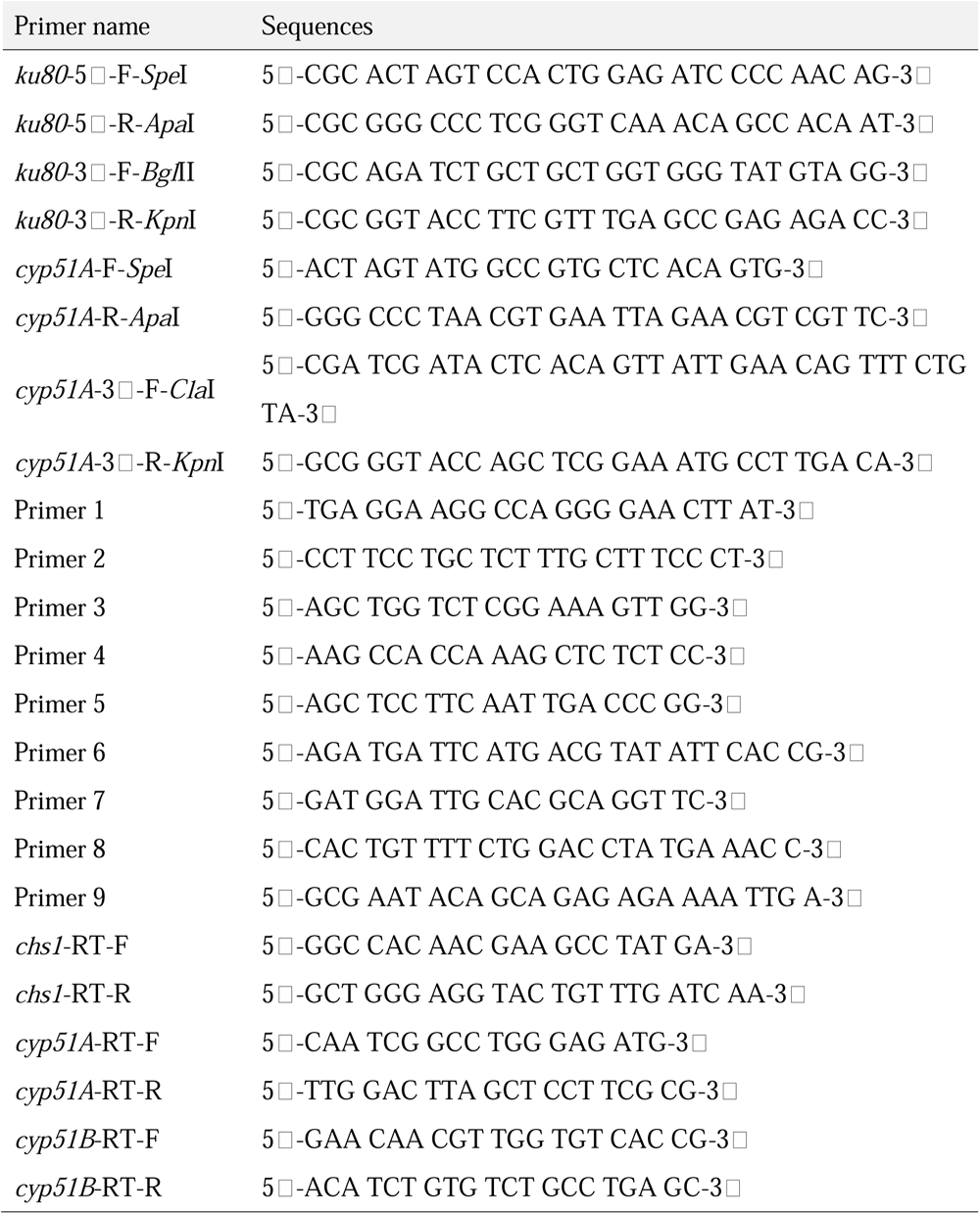
Primers used in this study.

### Transformation of *T. rubrum*

*T. rubrum* was transformed using the polyethylene glycol (PEG) method as described previously (12). The desired transformants and purified genomic DNA were analyzed by PCR. Total DNA was extracted using the Quick-DNA Fungal/Bacterial Miniprep Kit (Zymo Research, USA). Fungal cells were disrupted by μT-01 (TAITEC, Japan) using 5-mm stainless beads.

### Antifungal susceptibility assay

Conidia (2L × L10^3^) were incubated with two-fold serial dilutions of antifungal agents in 200 μl MOPS-buffered RPMI (pH 7.0) at 28°C for 7 days, and the minimum inhibitory concentration MIC_100_ (concentration required to inhibit growth by 100%) was determined. Efinaconazole was purchased from BLD Pharmatech Ltd, China, and ravuconazole was purchased from Merck.

### Quantitative reverse transcription-PCR (qRT-PCR)

Total RNAs were purified using NucleoSpin RNA (Macherey-Nagel) and reverse-transcribed into cDNAs using ReverTra Ace (Toyobo) according to the manufacturers’ instructions. qRT-PCR was performed using TB Green Premix Ex Taq II (TaKaRa Bio, Japan) on a StepOne Real-time PCR (Thermo Fisher Scientific, USA). The relative mRNA expression level was determined using the 2^-ΔΔ*Ct*^ method using *chitin synthase I* (*csh1*) as an endogenous control to normalize the samples (13). The primers used in this study are listed in Table 1.

### Statistical analysis

Mean values of three or more groups with two variables were compared using two-way ANOVA with Šidák correction (for Figure 2A) and Tukey’s post hoc test (for Figure 2D), according to the recommendation of Prism 10 (GraphPad, USA). The difference in the efficiency of HR in wild-type (WT) and Δ*ku80* strains was analyzed by two-sided Fisher’s exact test using Prism 10 (GraphPad, USA). Differences were considered significant at *P*L<L0.05.

## RESULTS

To increase gene targeting efficiency, we attempted to delete the gene encoding Ku80, which promotes nonhomologous recombination repair and whose deletion increases gene targeting efficiency by HR in other fungal species (7, 8). *T. rubrum* CBS 118892 has been isolated from human nail. This strain has been used for whole genome analysis (9) and several transcriptome analyses (14–17), as well as to produce genetically modified strains (18–21). Therefore, we used this strain as a parent strain of *ku80* deletion strain. The *ku80* ORF was replaced with a cassette with the neomycin resistance gene (*nptII*) and *flippase* (*flp*) flanked by flippase recognition sequences (**Fig. 1a**). As *flp* was inserted downstream of the copper ion-responsive promoter P*_ctr4_*, *nptII* and *flp* were removed from the *ku80*-deficient genome by adding the copper ion chelator bathocuproinedisulfonic acid to induce FLP recombinase expression (**Fig. 1a**).

**Figure 1.**
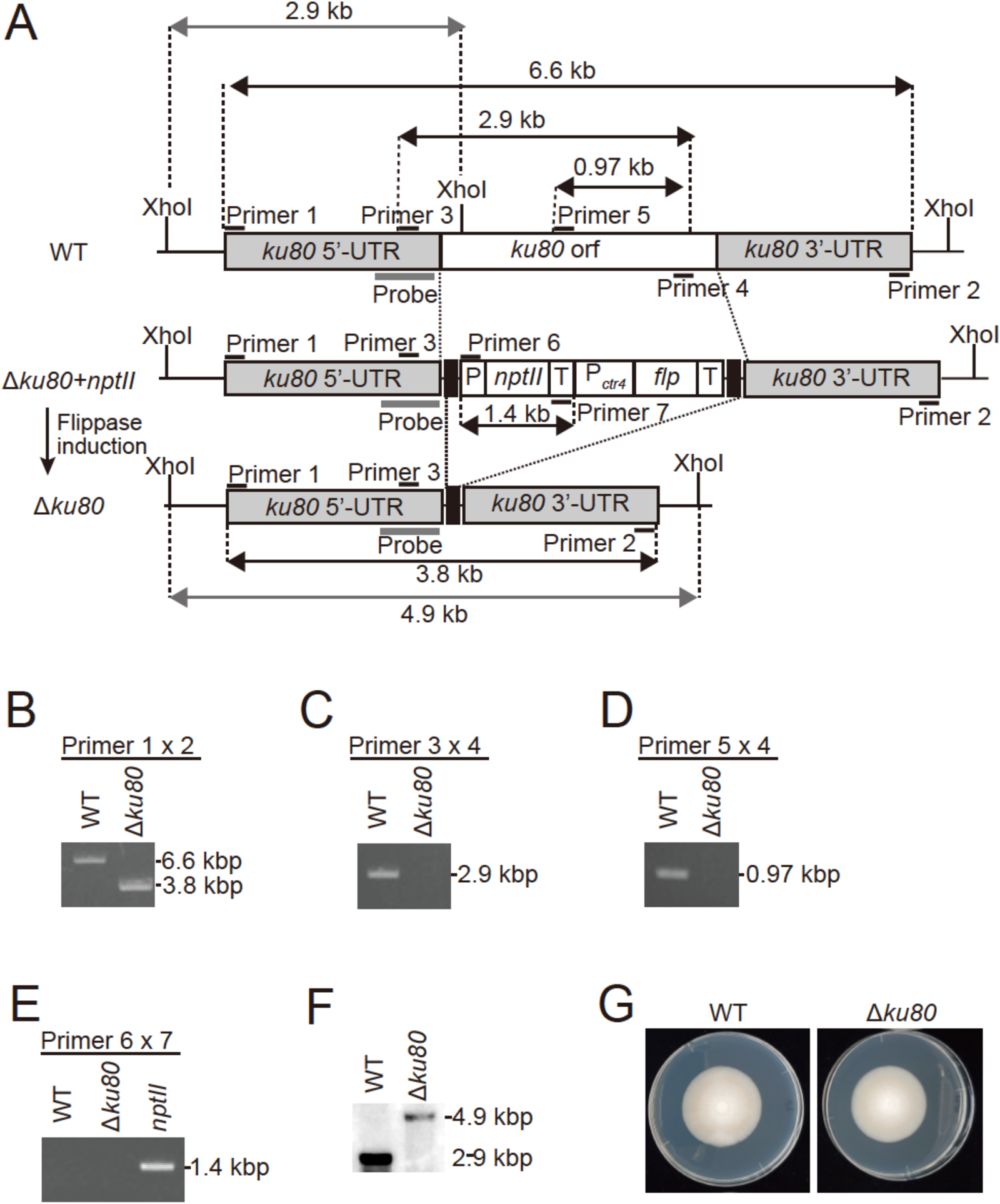
*ku80* locus targeting and *nptII* marker excision. (A) Schematic representation of the *ku80* locus before and after excision of flipper modules in *T. rubrum*. Site-specific recombination between the flanking FRT sequences (black box) was performed by the conditional expression of *flp*. (B–E) PCR analysis of total DNA samples from transformants. WT was used as a control. (B) Fragments were amplified with primer pairs (Primers 1 and 2). (C) Fragments were amplified with primer pairs (Primers 3 and 4). (D) Internal fragments of the *ku80* ORF were amplified with primer pairs (Primers 5 and 4). (E) Internal fragments of *nptII* were amplified with primer pairs (Primers 6 and 7). The *nptII*-harboring strain (Δ*cla4*) was used as a positive control. (F) Southern blot analysis of genome DNA samples from wild-type and Δ*ku80* strains. (G) Mycelial growth of WT and Δ*ku80* strains on SDA at 28°C for 16 days.

To confirm that the *ku80*-deficient strain (Δ*ku80*) was generated as designed, PCR was performed using genomic DNA purified from WT and Δ*ku80* strains (**Fig. 1a**, top and bottom, respectively) as templates. PCR performed using WT genomic DNA and primers designed for the 5L- and 3L-UTR of *ku80* (Primers 1 and 2 in **Fig. 1a**, respectively) amplified the PCR products with the expected size (6.6 kbp; **Fig. 1b**, left lane). The size of PCR products in Δ*ku80* was reduced as expected (3.8 kbp; **Fig. 1b**, right lane). In contrast, PCR performed using primers designed against sequences in the 5L-UTR (Primer 3 in **Fig. 1a**) and the ORF of *ku80* (Primers 4 and 5 in **Fig. 1a**) yielded PCR products of the expected size for WT (**Fig. 1c** and **d**, left lanes) but not for Δ*ku80* (**Fig. 1c** and **d**, right lanes) strain. The deletion of *nptII* from the genome of Δ*ku80 + nptII* strain (**Fig. 1a**, middle) was confirmed by PCR using primers designed against the sequences in the promoter and terminator of *nptII* (Primers 6 and 7 in **Fig. 1a**, respectively, **Fig. 1e**). The deletion of Δ*ku80* was also confirmed by Southern blot analysis of genomic DNA from WT and Δ*ku80* strains (**Fig. 1f**). These data indicated that the Δku80 strain was successfully generated with no reduction in the number of available drug markers. To ascertain the extent of the impact of Ku80 protein on growth, we compared mycelial growth between WT and Δ*ku80* strains, which revealed comparable mycelial growth (**Fig. 1g**).

Azole antifungal drugs used for treating dermatophytosis target the lanosterol demethylase Cyp51, which functions in the ergosterol synthesis pathway. XP_003235929 and XP_003236980 in *T. rubrum* have been identified as Cyp51A and Cyp51B homologs, respectively (22). It has been reported that the addition of azole antifungal drugs induces fungal Cyp51 expression (23, 24). In this study, the mRNA expression of *cyp51A* in *T. rubrum* was upregulated by the addition of the azole antifungal drug efinaconazole, but that of *cyp51B* was not upregulated (**Fig. 2a**). This result suggests that Cyp51A functions as a responsible Cyp51 isozyme when ergosterol biosynthesis is hindered, such as during treatment with azole antifungals. Because the *cyp51* homolog *erg11* is an essential gene in budding yeast, a deficiency of dermatophyte *cyp51A* could cause strong growth defects. In budding yeast, disruption of the natural 3′-UTR by the insertion of an antibiotic-resistant marker was found to destabilize the corresponding mRNAs, and this strategy has been used to analyze essential genes (25, 26). We attempted to insert the *neomycin phosphotransferase* (*nptII*) gene cassette into the downstream of *cyp51A* ORF of *T. rubrum*, as demonstrated in budding yeast studies (25, 26). Using the obtained Δ*ku80* strain, we inserted a *nptII* cassette into *cyp51A* 3L-UTR (hereinafter termed the 3L-UTR disruptant; **Fig. 2b** and **c**). Homologous recombinant strains were obtained in 12 of 26 strains (46.2%; Table 2) in which the insertion of the drug resistance gene was confirmed by PCR using primers designed within the ORF and 3L-UTR of *cyp51A* (Primers 8 and 9, respectively; **Fig. 2b** and **c**). The HR efficiency of the Δ*ku80* strain was 46 times higher than that of the WT strain (1/98; 1.0%; Table 2). These data demonstrated that a highly efficient HR method had been established in *T. rubrum*.

**Figure 2.**
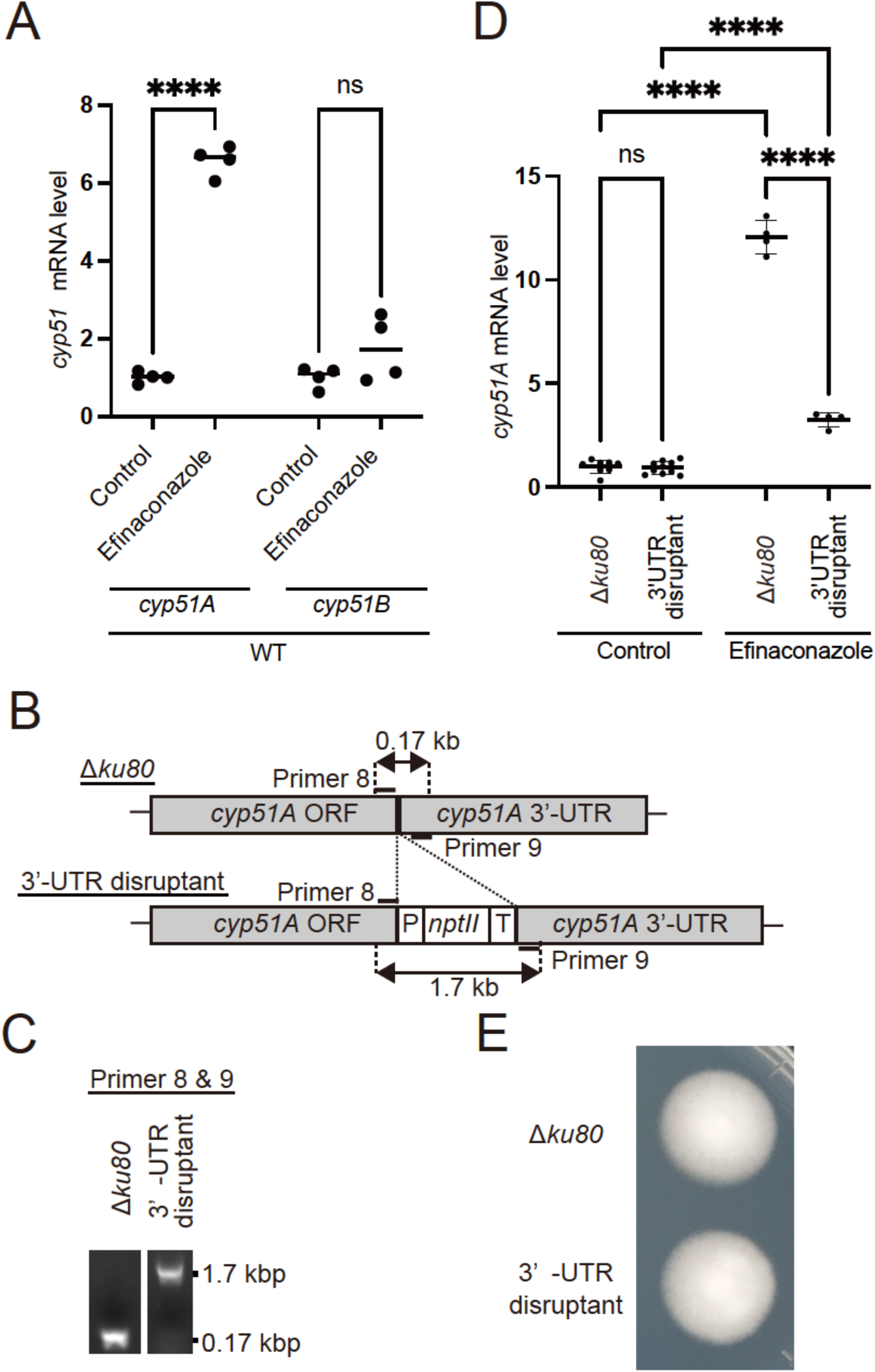
Production and characterization of the *cyp51A* 3**D**-UTR disruptant of *T. rubrum*. (A) The mRNA expression of *cyp51A* and *cyp51B* with or without 1/10 MIC of efinaconazole in WT. Data are expressed as mean ± SD. The dots on the graph represent biological replicates (n = 4). n.s., not significant. ****, P < 0.0001. (B) Schematic representation of the *cyp51A* locus of WT and *cyp51A* 3L-UTR disruptant (3L-UTR disruptant). (C) PCR analysis of total DNA samples from the 3L-UTR disruptant. The fragments were amplified with primer pairs (Primer 8 and 9). Δ*ku80* was used as a control. (D) The mRNA expression of *cyp51A* in Δ*ku80* and *cyp51A* 3L-UTR disruptant (3L-UTR) with or without 1/10 MIC of efinaconazole. The bars represent the standard deviation of the data obtained from three independent experiments. Data are expressed as mean ± SD. The dots on the graph represent biological replicates (n = 4–10). n.s., not significant. ****, PL<L0.0001. (E) Mycelial growth of Δ*ku80* and *cyp51A* 3L-UTR disruptant on SDA at 28L for 13 days.

**Table 2.** Gene targeting efficiency of WT and Δ*ku80* strains.

| Strain | Total transformants | Homologous replacement | Efficiency (%) |
| --- | --- | --- | --- |
| WT | 98 | 1 | 1.0 |
| $\Delta ku80$ | 26 | 12 | 46.2 |

\*Two-sided Fisher's exact test, $P < 0.0001$

The mRNA level of *cyp51A* in the 3′-UTR disruptant was comparable to that in the parent strain Δ*ku80* (**Fig. 2d**). Nevertheless, efinaconazole-induced elevation of *cyp51A* mRNA level decreased in the 3′-UTR disruptant (**Fig. 2d**). These findings suggest that the lack of the 3′-UTR causes mRNA perturbation at least under the condition of *cyp51A* induction in *T. rubrum*. Previous research has reported increased sensitivity to the azole antifungals fluconazole and itraconazole by the suppression of *cyp51A* expression in *A. fumigatus*, which has two Cyp51 isozymes, Cyp51A and Cyp51B, similar to that in *T. rubrum* (27). The 3L-UTR disruptant exhibited similar mycelial growth as that of the Δ*ku80* strain (**Fig. 2e**), but it showed increased sensitivity to the azole antifungals efinaconazole and ravuconazole (Table 3). However, the MICs of itraconazole and luliconazole remained unchanged. These findings suggest that Cyp51A functions as a factor for azole antifungal tolerance in *T. rubrum*.

**Table 3.** MIC values (μg/ml) of efinaconazole and ravuconazole in Δ*ku80* and *cyp51A* 3L-UTR disruptants (3L-UTR disruptants).

| Azole drugs | *Δku80* | 3'-UTR disruptant #1 |
| --- | --- | --- |
| Efinaconazole | 0.010 | 0.0025 |
| Ravuconazole | 0.040 | 0.020 |
| Itraconazole | 0.50 | 0.50 |
| Luliconazole | 0.00063 | 0.00063 |

## DISCUSSION

*T. rubrum* is an anthropophilic dermatophyte specialized for human parasitism, whereas several other dermatophytes are zoophilic or geophilic (28). The nature of this fungus is of great interest from not only a medical but also biological point of view. In recent years, transcriptomic, proteomic, and immunological studies of this fungus have been conducted extensively (17, 29–34). Nevertheless, molecular and cellular biological studies of *T. rubrum* have been limited partially due to a lack of genetic tools for this organism. In this study, we generated a *ku80*-deficient strain of this fungus and demonstrated that this strain can be applied in efficient HR methods, similar to a system established in a zoophilic dermatophyte, *T. mentagrophytes* (formerly *Arthroderma vanbreuseghemii*) (35). The method established in this study might serve as a fundamental technique to promote research that will advance the findings of previous comprehensive analyses and immunological analyses observed on the host side (17, 29–34).

The 3′-UTR disruptant, in which the expression induction of *cyp51A* by efinaconazole was attenuated, exhibited increased sensitivity to efinaconazole and ravuconazole. Considering that *cyp51A* expression was upregulated in response to efinaconazole addition, we speculated that *T. rubrum* Cyp51A is an inducible Cyp51 isozyme crucial for tolerance to azole antifungals. In *A. fumigatus*, loss or suppression of *cyp51A* expression enhances sensitivity to the azole antifungal fluconazole (27). Conversely, *cyp51B* deficiency does not significantly alter fluconazole sensitivity (27). This difference may be partially explained by the lower binding affinity of Cyp51A for fluconazole than for Cyp51B (36). Nevertheless, a difference in the induction of the expression of each *cyp51* gene in response to azoles may also contribute to this disparity in sensitivity. Regarding *T. rubrum*, no studies have investigated the contribution of Cyp51A and Cyp51B isozymes to the resistance to azole antifungal drugs. In the future, it is important to generate *T. rubrum* strains that are deficient in *cyp51A* and *cyp51B*, followed by analyzing their involvement in growth and resistance to azole antifungal drugs.

## Acknowledgments

The authors thank H. Uga, N. Hori, H. Hamanaka for their technical help. This work was supported by the Japan Society for the Promotion of Science.

## Author Contribution Statement

MI conceived and designed research. MI conducted experiments. MI, TY and SO analyzed data. MI, TY and SO wrote the manuscript. All authors read and approved the manuscript.

